# Evolution of the folding landscape of effector caspases

**DOI:** 10.1101/2021.08.05.455265

**Authors:** Suman Shrestha, A. Clay Clark

## Abstract

Caspases are a family of cysteinyl proteases that control programmed cell death and maintain homeostasis in multicellular organisms. The caspase family is an excellent model to study protein evolution because all caspases are produced as zymogens (procaspases) that must be activated to gain full activity; the protein structures are conserved through hundreds of millions of years of evolution; and some allosteric features arose with the early ancestor while others are more recent evolutionary events. The apoptotic caspases evolved from a common ancestor into two distinct subfamilies: monomers (initiator caspases) or dimers (effector caspases). Differences in activation mechanisms of the two subfamilies, and their oligomeric forms, play a central role in the regulation of apoptosis. Here, we examine changes in the folding landscape by characterizing human effector caspases and their common ancestor. The results show that the effector caspases unfold by a minimum three-state equilibrium model at pH 7.5, where the native dimer is in equilibrium with a partially folded monomeric (procaspase-7, common ancestor) or dimeric (procaspase-6) intermediate. In comparison, the unfolding pathway of procaspase-3 contains both oligomeric forms of the intermediate. Overall, the data show that the folding landscape was first established with the common ancestor and was then retained for >650 million years. Partially folded monomeric or dimeric intermediates in the ancestral ensemble provide mechanisms for evolutionary changes that affect stability of extant caspases. The conserved folding landscape allows for the fine-tuning of enzyme stability in a species-dependent manner while retaining the overall caspase-hemoglobinase fold.

## Introduction

While folding landscapes of proteins have been studied for decades (1–3), many studies focused on small monomeric proteins or dimers with simple folding landscapes (4, 5). Studies of monomeric proteins have provided a wealth of information concerning the principles that govern intramolecular interactions during folding, but they do not consider intermolecular interactions provided by the interfaces of multimeric proteins (6). For some dimers, subunit interactions in the dimer interface lead to more complicated folding pathways when compared to simple two-state behavior involving only native dimer (N_2_) and unfolded monomers (U) (7, 8). Moreover, relatively little is known about the evolution of dimeric proteins compared to monomeric proteins (2, 9), although two-thirds of proteins form a multimeric assembly (10, 11). Thus, an understanding of the role of oligomerization in the folding landscape of a polypeptide sequence should include a consideration of the interface in assembly (12). In this regard, the caspase family of proteases is an attractive model system to study the evolution of protein folding landscapes. Caspases are a family of cysteinyl aspartate-specific proteases that initiate and execute programmed cell death and maintain cellular homeostasis in metazoans (13). There are two broad categories of caspases based on their function: inflammatory caspases (caspases −1, −4, and −5) and apoptotic caspases (Fig. 1A). The latter is further subdivided into two groups based on their entry into the apoptotic cascade. Initiator caspases (caspases-2, −8, −9 and −10) act upstream and activate downstream effector procaspases (caspases-3, −6, and −7), which execute the cell death function (14). In addition, the caspase-hemoglobinase fold has been conserved for >650 million years (15), so the caspase family provides opportunities to examine the folding of monomers as well as changes to the folding landscape that resulted in oligomerization.

**Figure 1:**
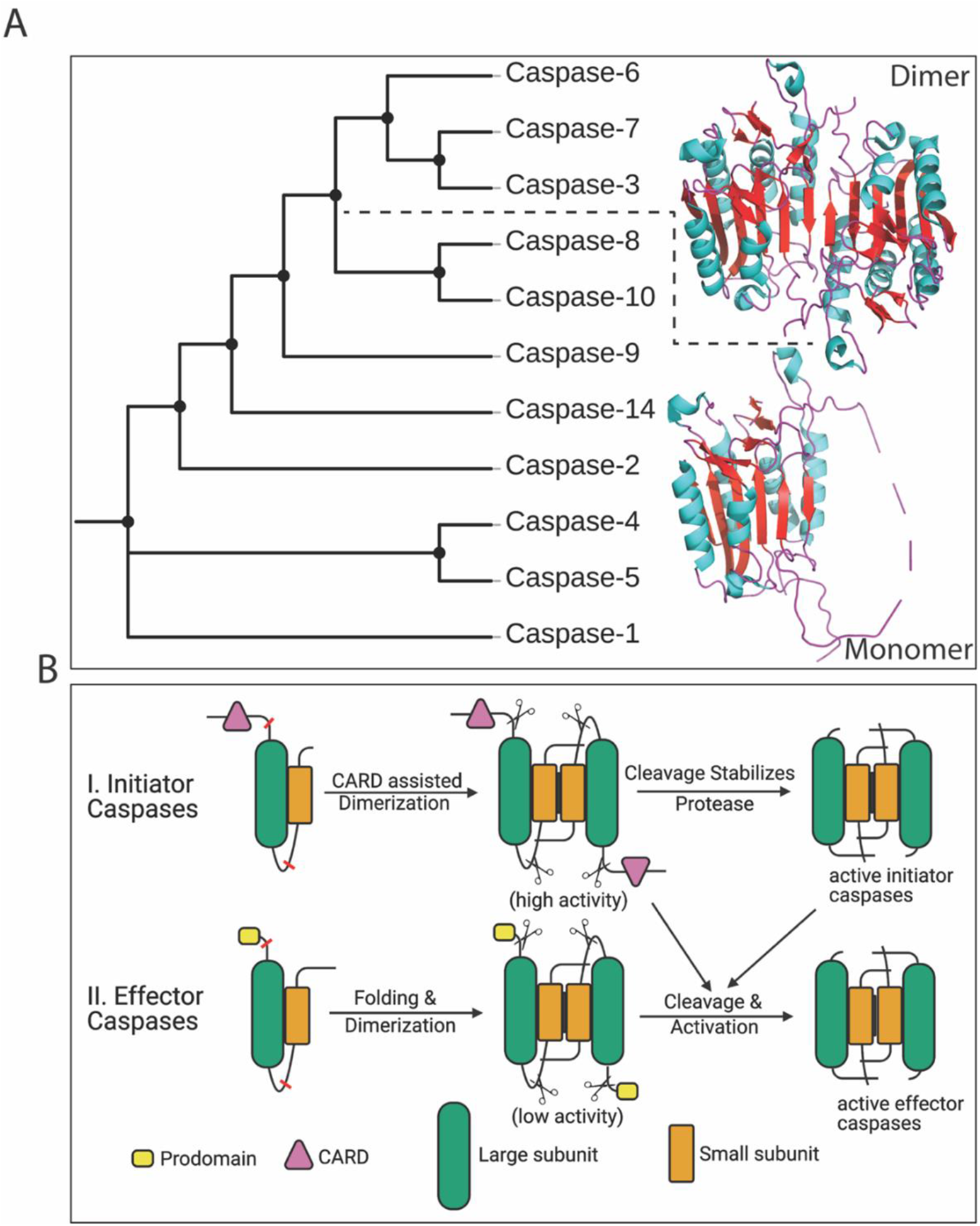
Phylogenetic relationship and activation mechanism of apoptotic caspases. (**A**) All caspases evolved from a common ancestor, then inflammatory (caspases-1, −4, −5) and initiator caspases (caspases-8, −9, −10, −14, −2) evolved as a monomer while effector caspases (caspases-3, −6, −7) evolved as dimers. (**B**) Initiator caspase zymogens are stable monomers, and dimerization is sufficient for activation, while effector caspases are stable dimers that are activated by cleavage of an intersubunit linker. Red lines represent cleavage sites and scissors represents cleavage.

More recently, evolutionary biochemists have used a new approach to study protein folding by examining the evolution of protein structure and function. Ancestral reconstruction methods enable one to use vertical comparisons (comparing ancestral to modern enzymes) as well as horizontal comparisons (comparing modern enzymes from multiple species) (16–18). The results of such studies potentially show the mechanisms by which changes in protein sequence have caused shifts in structure and function (19). The sequence determinants of protein structure-function, and substitutions revealed by the evolutionary analysis in common evolutionary nodes can then be introduced singly or in combination into ancestral backgrounds. Ultimately, by examining the ancestral reconstructions and changes that occur throughout evolution of the protein, one can determine the effects of historical mutations on protein structure, function, and physical properties (18).

The effector and initiator caspases evolved by gene duplication and divergence from a common ancestor (CA) more than 650 million years ago (Mya) (Fig. 1A) (15). The CA provided a scaffold for the modern caspases to acquire distinct properties, such as formation of oligomers, changes in stability, enzyme specificity, and allosteric regulation (15). All caspases are produced in the cell as inactive zymogens that must be activated during the apoptotic cascade. In general, initiator procaspases are stable monomers, and dimerization is sufficient for activation, whereas effector procaspases are stable, yet inactive dimers, and are activated via cleavage by initiator caspases (Fig. 1B). The oligomeric form of the zymogen and its activation mechanism is key to regulating apoptosis (13, 15). While the amino acid sequence identity is low between caspase subfamilies (~40%) (15), the caspase-hemoglobinase fold is well-conserved (20) even from distantly related species of vertebrates and invertebrates, such as human, *Danio rerio*, *Caenorhabditis elegans*, *Drosophila melanogaster,* and *Porites astreoides* (21–25).The structure of the procaspase monomer is characterized by a 6-stranded β-sheet core with several α-helices on the surface (6). Each monomer of the procaspase homodimer consists of approximately 300 amino acids organized into an N-terminal prodomain followed by a protease domain. The protease domain is further divided into large and small subunits that are connected by a short intersubunit linker. (Fig. 1B).

Previously, we showed that the human procaspase-3 dimer assembles by a four-state equilibrium mechanism in which the unfolded protein partially folds to a monomeric intermediate. Following dimerization of the intermediate, the protein undergoes a conformational change to form the native dimer (2U ⇄ 2I ⇄ I_2_ ⇄ N_2_) (26). Dimerization results in a substantial increase in conformational free energy, ΔG°_conf_, for the dimer (~25 kcal mol^−1^) *vs.* the monomer (~7 kcal mol^−1^) at 25°C (26). Furthermore, by examining the changes in ΔG°_conf_ *vs.* pH, we showed that the procaspase-3 dimer dissociates at lower pH due to a pair of histidine residues that contribute to salt bridges across the dimer interface (27). Thus, our previous data showed that the per residue contribution to the total conformational free energy of the dimer (ΔG°_conf_) increases from 0.025 kcal/mol/aa to 0.044 kcal/mol/aa (26, 27). However, because human procaspase-3 is the only caspase in which the folding properties have been examined, the evolutionary trajectories that resulted in dimer formation remain unknown.

We recently used ancestral protein reconstruction (APR) techniques to resurrect the common ancestor (CA) of the caspase-3/6/7 subfamily (28). In those studies, we examined the robustness of reconstruction methods by resurrecting two sequences (called AncCP-Ef1 and AncCP-Ef2) from the pool of possible sequences of CA, and we characterized the proteins biochemically and structurally (15). Here, we examine the evolution of the caspase folding landscape using the ancestral reconstruction AncCP-Ef2 (referred to here as PCP-CA, procaspase-common ancestor). The results are compared to those for extant human procaspase-6 (called PCP6) and procaspase-7 (called PCP7) as well as our previous studies for procaspase-3 (called PCP3) (26, 27). We examined urea-induced equilibrium unfolding over a broad pH range to compare the folding pathways of all three effector caspase subfamilies. The data show that the caspase folding landscape was first established and then retained for >650 million years. The common ancestor PCP-CA forms a weak dimer, and the dimer was stabilized early in the evolution of the subfamily. Of the three extant human effector caspases, caspase-6 is the most stable, while caspase-7 is the least stable. The folding landscape of procaspase-7 is more similar to that of the common ancestor, which is consistent with previous phylogenetic data that show caspase-7 is closest to the common ancestor (15). In procaspases-3 and −6, folding intermediates were stabilized later in evolution.

## Results

A protomer of PCP6, PCP7, and PCP-CA consists of 293, 303, and 275 amino acids, respectively (Supplemental Table S1). Under some conditions, such as the high protein concentrations found in heterologous expression systems, effector caspase activation is autocatalytic. In order to prevent autoproteolysis during expression in *E. coli*, we substituted the active site cysteine with a serine residue for our equilibrium unfolding studies, as described previously for procaspase-3 (26).

PCP6 and PCP7 have two tryptophan residues, while PCP-CA has only one tryptophan (Fig. 2 & Supplemental Table S1). In PCP6, one tryptophan resides in active site loop 1 (Supplemental Fig. S1), whereas the second tryptophan resides in active site loop 3. In PCP7 and in PCP-CA, the tryptophans are found only in active site loop 3 (Fig. 2 and Supplemental Fig. S1). PCP6, PCP7, and PCP-CA have 10, 9, and 13 tyrosine residues, respectively, and they are well distributed in the primary sequence (Fig. 2 and Supplemental Fig. S1). Native PCP6 has a fluorescence emission maximum at 319 nm (Fig. 3A and 3B), while that of native PCP7 is at 338 nm, when excited either at 280 nm or 295 nm (Fig. 3C and 3D). In the case of an PCP-CA, the fluorescence emission maximum is 321 nm when excited at 280 nm and 330 nm when excited at 295 nm (Fig. 3E and 3F). Overall, the data show that the tryptophan residues in PCP7 are more solvent-exposed than those of PCP6 or of PCP-CA. In phosphate buffer containing 8 M urea, the fluorescence emission maximum is red shifted to ∼350 nm following excitation at 280 nm or 295 nm in all three proteins, indicating that the proteins were largely unfolded under these solution conditions (Fig. 3). At intermediate concentrations of urea (3 to 5 M), the emission maxima were red-shifted in the case of PCP6 and PCP-CA, but were largely unaffected in the case of PCP7. The changes in emission maxima are described more fully below.

**Figure 2:**
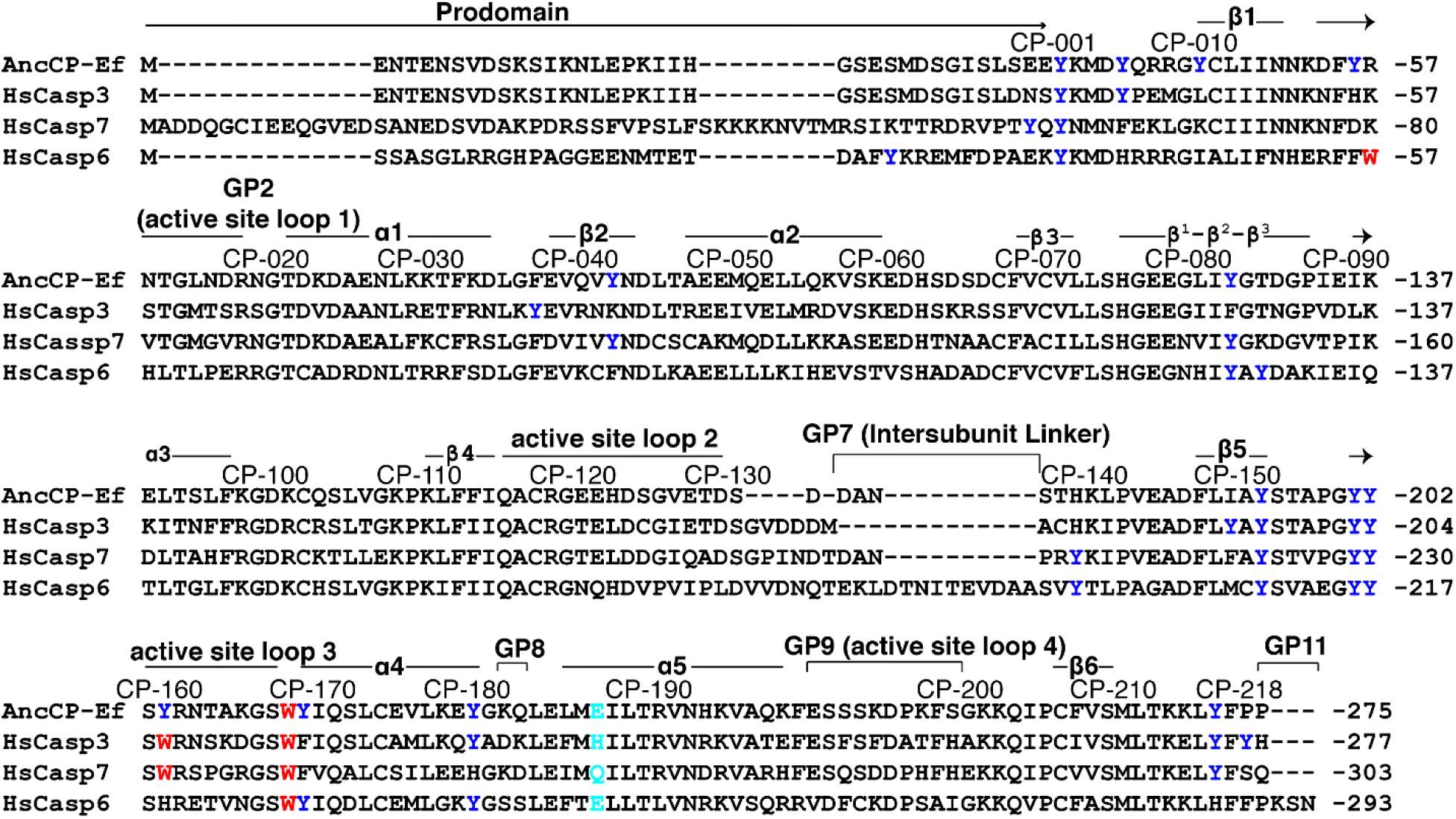
Multiple sequence alignment of human effector caspases with a common ancestor. CP refers to common position number of caspases (52), individual sequence number is indicated at the right side of sequences. Secondary structural elements are indicated, and tyrosine residues (blue), tryptophan residues (red), and CP-186 (cyan), unique amino acid among effector caspases in dimeric interface are highlighted.

**Figure 3:**
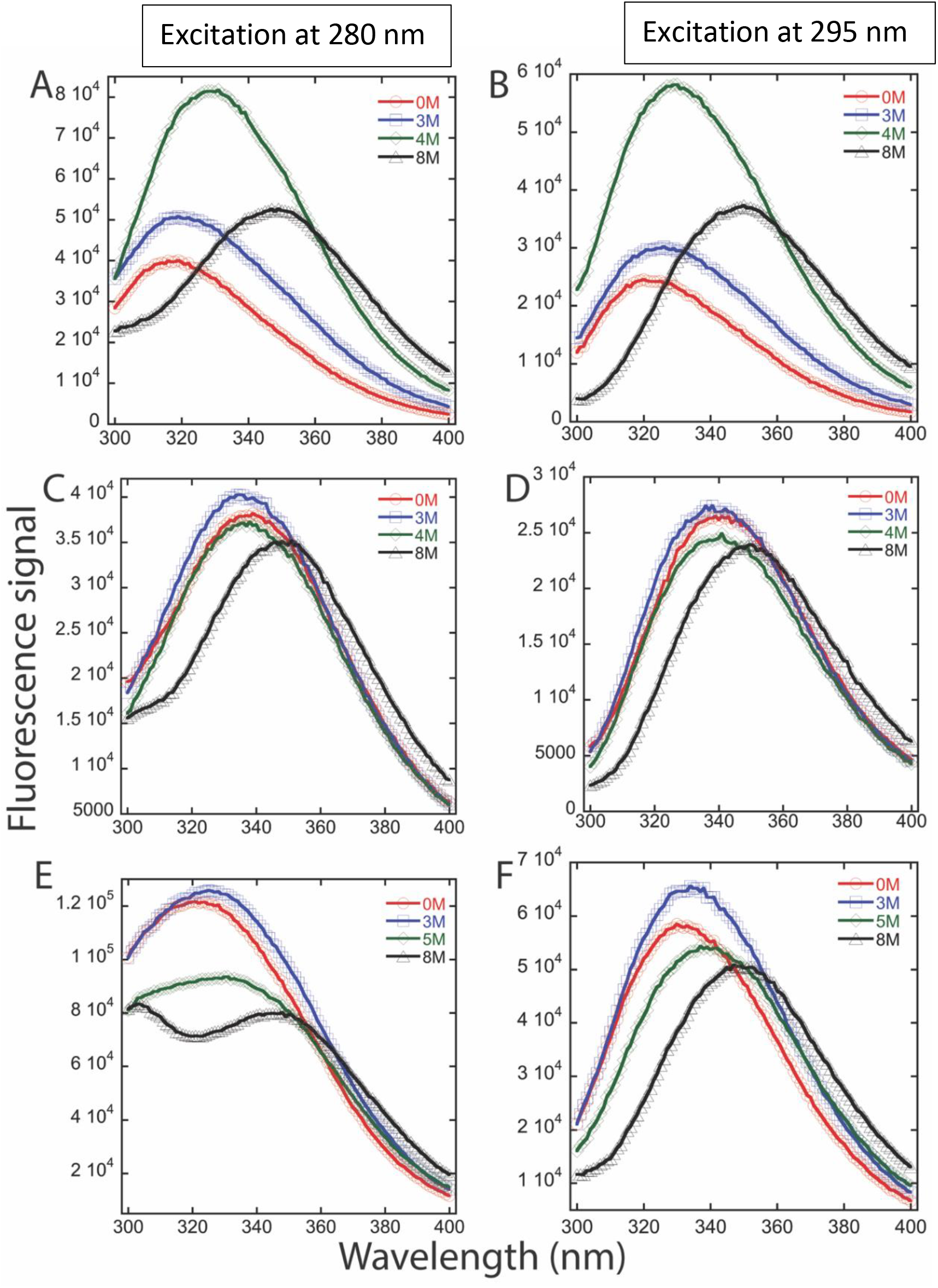
Fluorescence emission spectra of effector caspases following excitation at 280 nm (left) or 295 nm (right). Emission spectra of PCP6 (**A, B**), PCP7 (**C, D**), and PCP-CA (**E, F**) at 2 μM protein concentration in a buffer of 20 mM phosphate, pH 7.5, containing urea (displayed in inset of each graph).

### Equilibrium unfolding of extant and ancestor caspases

Changes in the fluorescence emission and circular dichroism properties of PCP3 as a function of urea concentration have been described previously (26, 27). In this study, we examined the equilibrium unfolding of PCP6, PCP7, and PCP-CA at pH 7.5 as a function of urea concentration (0 - 8 M), and the results are shown in Figure 4. Renaturation experiments of all three proteins demonstrated the folding transitions are reversible.

**Figure 4:**
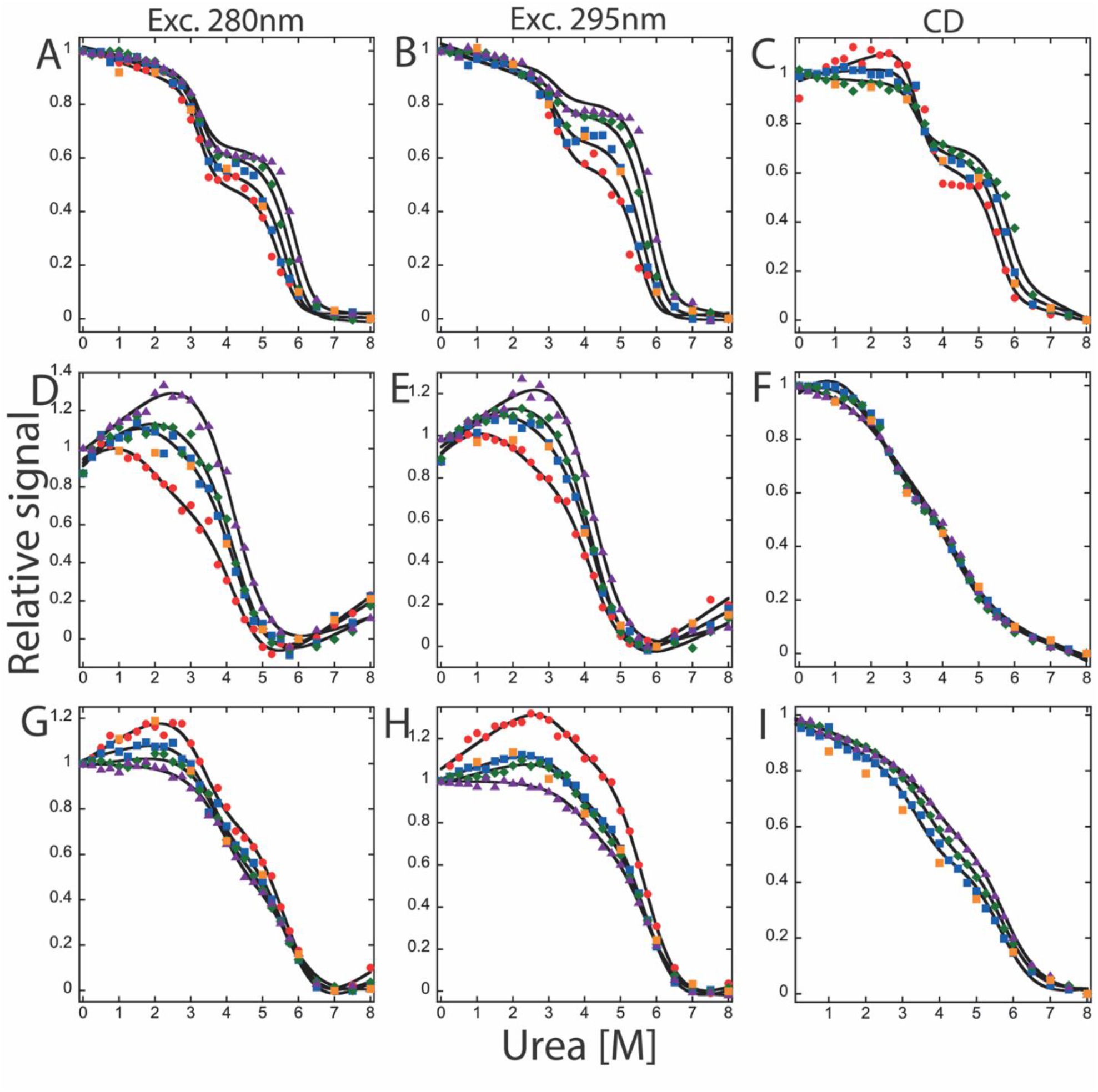
Equilibrium unfolding of effector caspases at pH 7.5. Equilibrium unfolding of PCP6 (**A, B, C**), PCP7 (**D, E, F**), and PCP-CA (**G, H, I**) monitored by fluorescence emission with excitation at 280 nm (left column), 295 nm (middle column) and circular dichroism (right column). Four different protein concentrations were used to measure unfolding monitored by fluorescence emission, and three different protein concentrations were used in to monitor unfolding by CD. Colored solid symbols represent raw data and corresponding solid lines represent the global fits of the data to an appropriate model as described in text. The following protein concentrations were used: 0.5 μM (●), 1 μM (◼), 2 μM (◆), and 4 μM (▲). Orange squares (◼) represent refolding data of 1uM protein to show reversibility.

For PCP6 at pH 7.5, both the fluorescence emission data (Fig. 4A and 4B) as well as the CD data (Fig. 4C) show little to no change in signal between 0 and ∼2.5 M urea. One then observes a cooperative decrease in the signal between ∼3.5 and 5 M urea, demonstrating a plateau between the native and unfolded signals. A second cooperative transition occurs between ∼5 and 6.5 M urea. For PCP6, the relative signal of the plateau as well as the second cooperative transition are dependent on the protein concentration.

In contrast to PCP6, both PCP7 (Fig. 4D-4F) and PCP-CA (Fig. 4G-4I) show a protein-concentration dependent increase in the relative signal between 0 and ∼3 M urea, followed by a cooperative decrease in the signal between ~3 and 6 M urea to form the unfolded state. Thus, the unfolding data are similar for PCP7 and PCP-CA, where the protein concentration dependence is observed in the first unfolding transition rather than the second transition, as in the case of PCP6.

Altogether, the data in Figures 3 and 4 show that the three proteins fold through a stable intermediate, which can be characterized by a red-shift in fluorescence emission and a concomitant loss of secondary structure compared to the native dimer. In the case of PCP6, however, the protein concentration dependence is observed in the second transition, whereas both PCP7 and PCP-CA show a protein concentration dependence in the first transition. Overall, the data suggest that while the partially folded intermediates that form during urea-induced unfolding have more solvent-exposed tryptophans and less secondary structure, compared to the native dimer, the intermediate remains dimeric in the case of PCP6 while the intermediate is monomeric in the cases of PCP7 and PCP-CA.

### Global fitting of equilibrium unfolding data

The experimental design described above at pH 7.5 for monitoring fluorescence emission (three to four protein concentrations, each with two excitation wavelengths) and circular dichroism (three protein concentrations) provides eleven data sets for each protein are fit globally to determine the free energy and the cooperativity index (m-value) for each unfolding transition. In the case of PCP6, the data were best fit to the three-state equilibrium model described in equation 1. In this model, the dimeric native conformation, N_2_, isomerizes to a dimeric intermediate, I_2_, and the dimeric intermediate dissociates and unfolds to monomers. The dissociation of I_2_ to 2U leads to a protein-concentration dependent change in the mid-point of the second transition, as shown in Figures 4A - 4C. Based on this model, we have determined the conformational free energy, ΔG°_conf_, and the m-values for each step of unfolding. The solid lines in Figures 4A - 4C are the results of global fits of the model to the data. The free energy change, 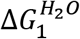 and the cooperativity index, m_1_, for the first step of unfolding, the isomerization of N_2_ to I_2_, are 8.4 ± 0.8 kcal/mol and 2.6 ± 0.3 kcal mol^−1^ M^−1^, respectively (Table 1). The free energy change, 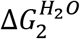, and cooperativity index, m_2_, for the dissociation and complete unfolding of the dimeric intermediate to two unfolded monomers (I_2_ ⇄ 2U) are 24.4 ± 0.9 kcal/mol and 2.9 ± 0.2 kcal mol^−1^ M^−1^, respectively. Overall, the data demonstrate that PCP6 is very stable, with the total conformational free energy of 32.8 kcal/mol at pH 7.5 and 25 °C.

**Table 1.** Thermodynamic parameters of each step of folding/unfolding of extant and ancestral effector caspases at pH 7.5.

| Proteins | Equilibrium mechanism | Free energy changes ($\Delta G^{\circ}_{\text{conf}}$) (kcal mol<sup>-1</sup>) | Cooperativity index (m-value) (kcal mol<sup>-1</sup> M<sup>-1</sup>) | Total $\Delta G^{\circ}_{\text{conf}}$ (kcal mol<sup>-1</sup>) | $m_{\text{total}}$ (kcal mol<sup>-1</sup> M<sup>-1</sup>) |
| --- | --- | --- | --- | --- | --- |
| PCP3* | $N_2 \rightleftharpoons I_2$ | $7.9 \pm 1.3$ | $2.8 \pm 0.5$ | $24.8 \pm 0.9$ | $4.5 \pm 0.7$ |
| | $I_2 \rightleftharpoons 2I$ | $9.7 \pm 1.0$ | $0.5 \pm 0.1$ | | |
| | $2I \rightleftharpoons 2U$ | $7.2 \pm 0.5$ | $1.2 \pm 0.1$ | | |
| PCP6 | $N_2 \rightleftharpoons I_2$ | $8.4 \pm 0.8$ | $2.6 \pm 0.3$ | $32.8 \pm 1.5$ | $5.5 \pm 0.5$ |
| | $I_2 \rightleftharpoons 2U$ | $24.4 \pm 0.9$ | $2.9 \pm 0.2$ | | |
| PCP7 | $N_2 \rightleftharpoons 2I$ | $10.2 \pm 0.2$ | $1.3 \pm 0.1$ | $15.4 \pm 0.3$ | $2.5 \pm 0.2$ |
| | $2I \rightleftharpoons 2U$ | $5.2 \pm 0.1$ | $1.2 \pm 0.1$ | | |
| PCP-CA | $N_2 \rightleftharpoons 2I$ | $14.9 \pm 0.1$ | $2 \pm 0.1$ | $21.8 \pm 1.3$ | $3.2 \pm 0.2$ |
| | $2I \rightleftharpoons 2U$ | $6.9 \pm 0.3$ | $1.2 \pm 0.1$ | | |

\* Data from Bose and Clark (26).

For PCP7 and PCP-CA, the global fits demonstrate that the data are well described by a three-state equilibrium model. In contrast to PCP6, described above, the partially folded intermediate is monomeric for PCP7 and for PCP-CA (equation 2). The solid lines in Figures 4D–4F (PCP7) and 4G–4I (PCP-CA) are the results of fits of the data to the model. In the case of PCP7, the free energy change, 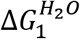, and the cooperativity index, m_1_, for the first step of unfolding, the dissociation of N_2_ to 2I, are 10.2 ± 0.2 kcal/mol and 1.3 ± 0.1 kcal mol^−1^ M^−1^, respectively. The free energy change, 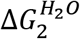, and cooperativity index, m_2_, for the complete unfolding of the monomeric intermediate to unfolded monomeric proteins (I ⇄ U) are 5.2 ± 0.1 kcal/mol and 1.2 ± 0.1 kcal mol^−1^ M^−1^, respectively (Table 1). Similarly, for PCP-CA, 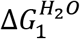, and m1 are 14.9 ± 0.1 kcal/mol and 2.0 ± 0.1 kcal mol^−1^ M^−1^, respectively. For unfolding of the monomeric intermediate of PCP-CA (I ⇄ U), 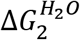 and m_2_, are 6.9 ± 0.3 kcal/mol and 1.2 ± 0.1 kcal mol^−1^ M^−1^, respectively (Table 1).

Overall, the data suggest a minimum three-state process in all effector caspases in which a well-populated intermediate is in equilibrium with the native and unfolded protein. Comparatively, of the three human effector caspases and the common ancestor, PCP6 is the most stable with ΔG°_conf_ of 32.8 kcal/mol, PCP-CA and PCP3 are intermediate with ΔG°_conf_ of 21.8 kcal/mol or 24.8 kcal/mol, respectively, and PCP7 is the least stable with ΔG°_conf_ of 15.4 kcal/mol at pH 7.5 (Fig. 5).

**Figure 5:**
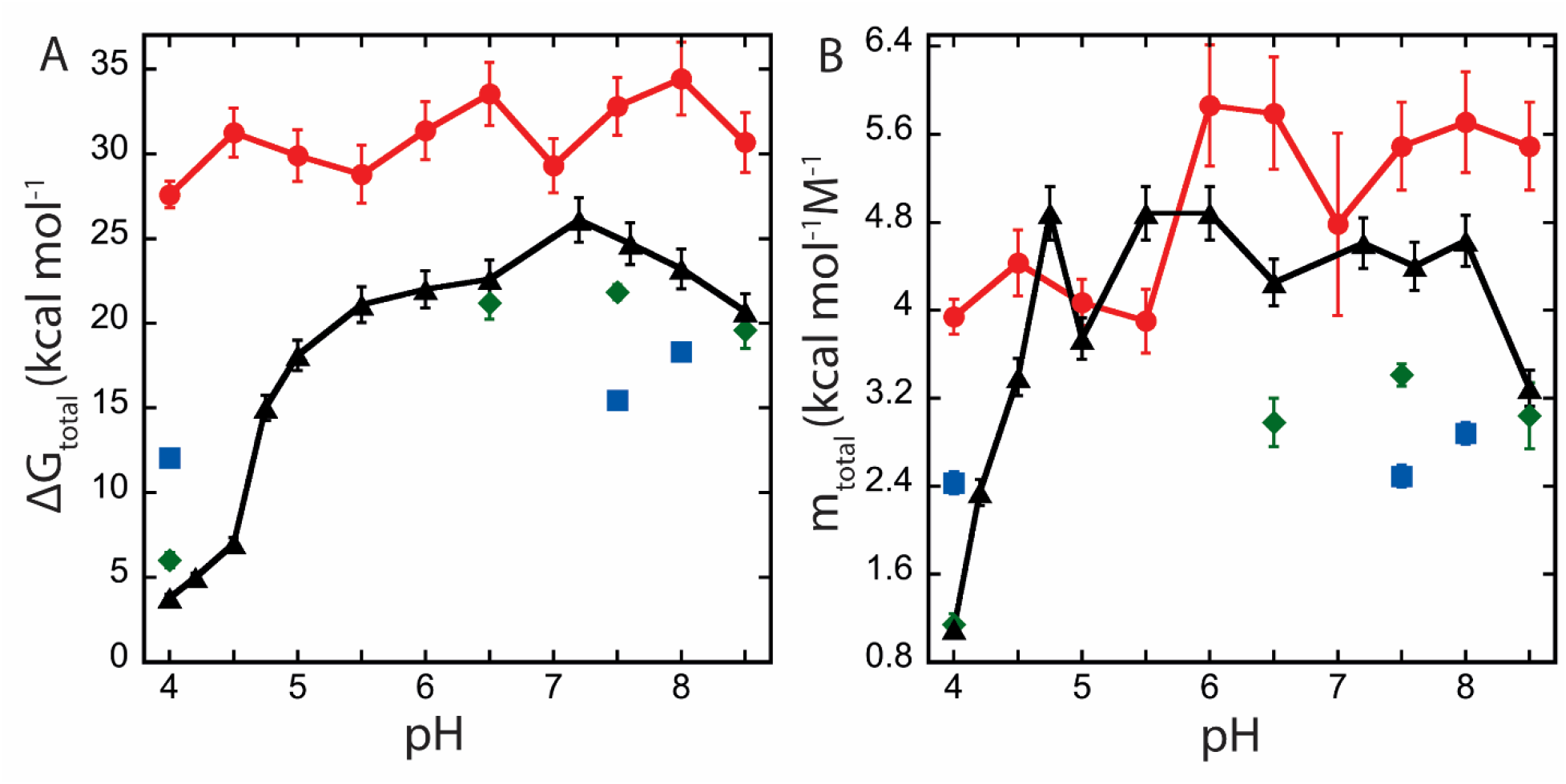
Comparison of total conformational free energy (ΔG_total_) (**A**), and total m-value (m_total_) (**B**) changes as a function of pH between PCP6 (●), PCP7 (◼), PCP-CA (◆), and PCP3 (▲). Error bars represent standard deviation of respective parameters determined by the global fitting. Data for PCP3 are from (27).

### pH effects on equilibrium unfolding of effector caspases

We showed previously that PCP3 undergoes pH dependent conformational changes, and the dimer dissociates below pH 5.5 (27). From our previous data, we suggested that dimer dissociation was due to a series of salt bridges across the dimer interface, which include two histidine residues. The other effector caspases also have charged amino acids that interact across the dimer interface, but only PCP3 contains the histidine residues. In order to determine the effects of pH on dimer stability, we performed equilibrium unfolding studies of PCP6, PCP7 and PCP-CA between pH 8.5 and pH 4. The data are summarized in Figure 5, and all folding/unfolding data are shown in Supplemental Figures S2-S6, while the conformational stability and m-value for each folding transition are shown in Supplemental Tables S2 and S3.

Similarly to the data described above at pH 7.5, the equilibrium folding/unfolding of PCP6 can be described by a three-state equilibrium model (eq. 1) over the pH range of 4.5 to 8.5. At pH 4, however, the data were best described by a two-state equilibrium model where the native dimer is in equilibrium with the unfolded monomer (eq 3) (Supplemental Figs. S2 and S3). Surprisingly, even at pH 4, PCP6 remains in a dimeric form, as demonstrated by the protein-concentration dependence to unfolding (Supplemental Figs. S2-S4). The protein stability, ΔG°_conf_, is ~30 kcal/mol throughout the entire pH range (Fig. 5), demonstrating the consistently high conformational free energy of the PCP6 dimer over a broad pH range. Based on the fits of the data to the corresponding equilibrium folding model (Supplemental Tables S2 and S3), we calculated the fraction of species *versus* urea concentration at each pH examined (Figure 6 and Supplemental Figure S4). The data show that the dissociation of the dimer is relatively consistent, with urea½ ~6 M throughout the entire pH range. In contrast, the fraction of native dimer, N_2_, decreases below pH 6 relative to the fraction of dimeric intermediate, I_2_, such that at pH 4 the dimeric intermediate is fully populated in the absence of urea (Supplemental Figure S4).

**Figure 6:**
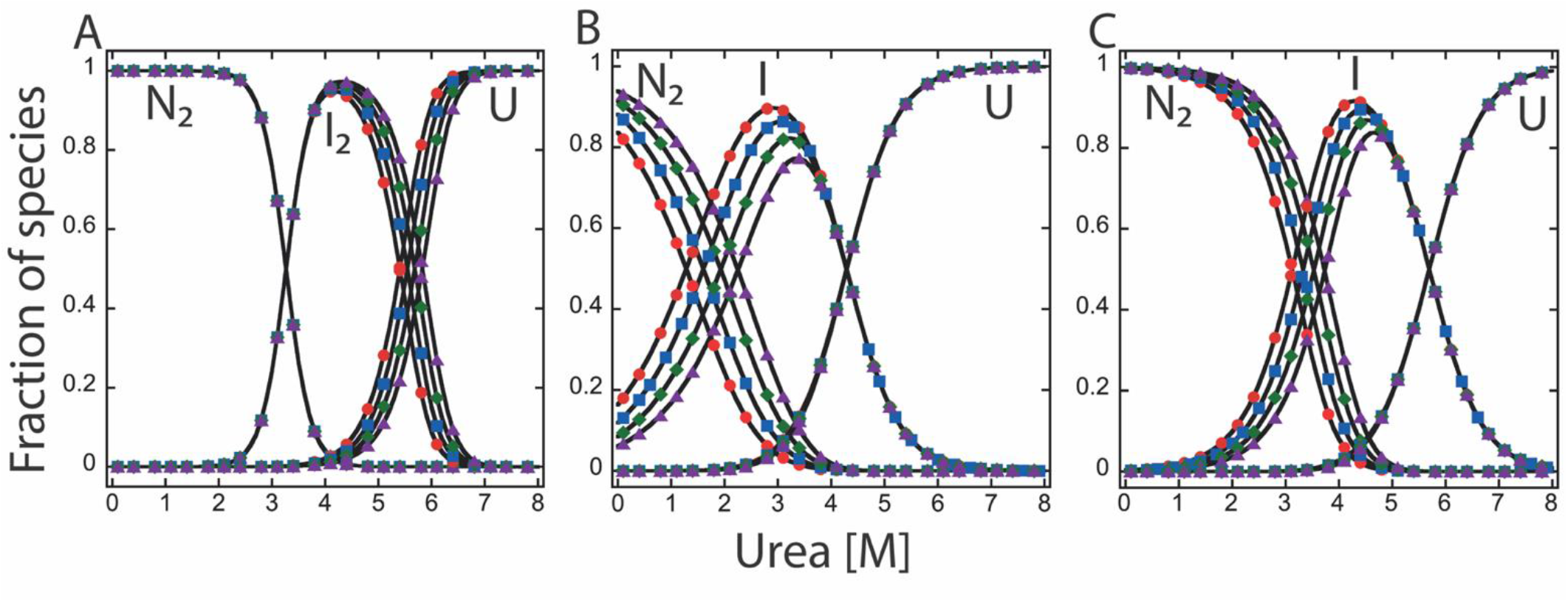
Fraction of species as a function of urea concentration at pH 7.5. Fraction of species of PCP6 (**A**), PCP7 (**B**), and PCP-CA (**C**). The fractions of native, intermediate, and unfolded protein were calculated as a function of urea concentration from fits of the data shown in Figure 4. N_2_ refers to dimeric native protein, I_2_ and I are dimeric and monomeric intermediates, respectively, and U refers to unfolded species. The following protein concentrations were used: 0.5 μM (●), 1 μM (◼), 2 μM (◆), and 4 μM (▲).

In the case of PCP7, the equilibrium folding/unfolding data were well described by the three-state equilibrium model, discussed above for data at pH 7.5, in which the native dimer is in equilibrium with a monomeric intermediate (eq. 2) (Supplemental Fig. S5). The conformational free energy (ΔG°_conf_) was 15-18 kcal/mol at higher pH (Supplemental Tables S2 and S3), similar to the data described above for pH 7.5. In contrast to PCP6, however, we were unable to examine the equilibrium folding/unfolding of PCP7 from pH 4.5 to pH 7 because protein aggregation resulted in irreversible unfolding. At pH 4, unfolding for PCP7 is reversible, and the data were also best fit to the three-state equilibrium model with a monomeric intermediate (eq. 2). The data suggest that the primary differences in the fluorescence emission at pH 8 versus pH 4 is that the monomeric intermediate exhibits a higher fluorescence emission relative to the native conformation at pH 8. At pH 4, however, the fluorescence emission of the intermediate is lower than that of the native conformation. Overall, the data show that, while the overall conformational stability of PCP7 is lower than that of PCP6, at all pHs, the unfolding of the monomeric intermediate has a similar urea½ of ~4.5M. In contrast, the native dimer is less stable at lower pH, although the protein appears to remain dimeric, and one observes that ΔG°_conf_ decreases by ~5 kcal/mol due to the lower dimer stability (Supplemental Table S2).

Similar to PCP7, we observed that PCP-CA does not fold reversibly from pH 4.5 to pH 6 due to aggregation. The data for PCP-CA at pH 6.5 to pH 8.5 were best described by a three-state equilibrium model in which a monomeric intermediate is in equilibrium with the native dimer and unfolded monomer (eq. 2) (Fig. 4G-4I and Supplemental Fig. S6), like PCP7. One observes a protein concentration dependence to unfolding between ~2M to 5M urea followed by a protein concentration-independent transition between ~5M to 7M urea. Based on global fits of the data to the three-state equilibrium model (eq. 2), we determined the total conformational free energy on unfolding of PCP-CA from pH 6.5 to pH 8.5 is ∼20 kcal/mol (Fig. 5, Table 1, and Supplemental Tables S2 and S3). In contrast, the equilibrium folding/unfolding data at pH 4 were the best fit to a three-state model in which the native monomer unfolds through a monomeric intermediate (eq. 4). We note that we cannot rule out that a small population of dimer is present at pH 4, but fitting the data to include a native dimer (eq. 2), dimeric intermediate (eq. 1), or no intermediate did not improve the quality of the fits. Our interpretation of the results is that the ΔG°_conf_ of the native species at pH 4 is very low (0.4 kcal/mol, see Supplemental Table S2), so it is difficult to observe a protein-concentration dependence to unfolding. A comparison of the changes in secondary structure observed by circular dichroism effectively illustrates the point, when compared at pH 8 versus pH 4 (compare Supplemental Figs. S6C and S6G). Thus, we selected the simplest model to describe the data, equation 4, which suggests that PCP-CA is a monomer at pH 4 and unfolds through a partially folded monomeric intermediate. In this case, the ΔG°_conf_ of unfolding is 5.5 kcal/mol (Supplemental Fig. S6 and Supplemental Tables S2 and S3).

The total conformational free energies of unfolding, ΔG°_conf_, and m-values obtained from the fits of all three proteins over the pH range of 4 to 8.5 are shown in Figures 5A and 5B, respectively. Due to aggregation upon refolding of PCP-CA at pH range 4.5 to 6, we were unable to determine the precise pH range for the transition of dimer to monomer. Nevertheless, PCP-CA is similar to PCP3 (27) in that the dimer is destabilized at lower pH, such that the protein is a monomer at pH 4. Together, the data show that the dimer is destabilized in PCP3 and PCP7 at lower pH, although PCP7, like PCP6, remains dimeric. Because the dimer contributes a substantial portion of the conformational free energy, PCP6 and PCP7 exhibit a relatively consistent ΔG°_conf_ at all pHs, while the ΔG°_conf_ of PCP3 and of PCP-CA reflect the stability of the monomer at pH 4 (Fig. 5). Unlike PCP3, however, we were unable to determine the pKa for dimer dissociation of PCP-CA due to protein aggregation between pH 4.5 and 6.5.

From the global fitting, we also calculated the cooperativity index (m-value) for each unfolding step. The m-value relates to the accessible surface area (ΔASA) exposed to solvent during unfolding (29). The total m-value (m_total_) of PCP6, PCP7 and PCP-CA range from 2.5-5.5 kcal mol^−1^ M^−1^ respectively at higher pH (Figure 5B, Table 1, and Supplemental Table S3). Scholtz and co-workers (29) developed an empirical analysis to show the correlation of ΔASA with experimental m-values (eq. 5). Using their analysis, our data suggest that the native dimer of PCP7 and of PCP-CA are less compact than those of PCP6 or PCP3. One observes that the m-values of the PCP7 and PCP-CA monomers are similar to that determined previously for PCP3 (~1.2 kcal mol^−1^ M^−1^) (Supplemental Table S3), suggesting that the surface area exposed during unfolding of the monomer is similar for the effector caspases. In contrast, the native dimer of PCP7 and of PCP-CA appear to have larger exposed surface area compared to those of PCP3 or of PCP6, resulting in a lower m-value (and related ΔASA) during unfolding of the dimer.

### The fraction of species versus urea concentration

For each pH, we calculated the equilibrium distribution of species over the urea concentration range of 0 to 8 M using the values obtained from the global fits of the equilibrium unfolding data (described above), the cooperativity indices determined for each transition (Supplemental Tables S2 and S3), and four protein concentrations (0.5, 1, 2, and 4 μM). The fraction of species for PCP6, PCP7, and PCP-CA are shown in Figure 6 and Supplemental Figures S4-S6. Collectively, at pH 7.5 one observes a cooperative decrease in native dimer (N_2_) with a concomitant increase in a partially folded intermediate between 0 and ∼4 M urea (Fig. 6). In the case of PCP6 and of PCP3 (27), the intermediate is dimeric (I_2_), while for PCP7 and PCP-CA the intermediate is a monomer (I). The dimeric (PCP6 or PCP3) or monomeric (PCP7 or PCP-CA) intermediate reaches a maximum at ∼3 to 4 M urea and remains predominant to ∼5 to 6 M urea (Fig. 6 and Supplemental Figs. S4 – S6). The unfolded state is fully populated by ~7 M urea in all cases. At pH 4, the “native” ensemble of PCP6 and of PCP7 consists of a dimeric conformation (Supplemental Figs. S4A and S5G), while the major fraction of PCP-CA (Supplemental Fig. S6D) consists of monomers. Together, the pH studies suggest that the native dimer (N_2_) of the effector caspases (PCP6, PCP3, PCP7) and of the common ancestor (PCP-CA) is destabilized at low pH relative to a partially folded intermediate, either the dimer, I_2_ (PCP6), or the monomer, I (PCP7, PCP3, PCP-CA).

Previously, we showed that the dimer of PCP3 undergoes a pH-dependent conformational change and that the pKa for the transition is ~5.7 (30). Likewise, the data for PCP6 suggests that a similar transition occurs in the dimer, with a similar pKa of ~5.9 (Supplemental Fig. S4). In this case, we examined the mid-point of the first transition at each pH (N_2_ ⇄ I_2_). In PCP3, however, a second transition occurs as the pH is lowered such the dimer dissociates, with a pKa ~4.7. Thus, PCP6 and PCP3 undergo a similar pH-dependent transition of N_2_ to I_2_, with pKa~5.7-5.9, but PCP6 remains dimeric at lower pH whereas the dimer of PCP3 dissociates.

## Discussion

The caspase-hemoglobinase fold is an ancient protein fold that has been conserved for at least 650 million years (15) and from which evolved three subfamilies of caspases (13). All caspases are produced in the cell as inactive zymogens that must be activated prior to their function in the inflammatory response or in apoptosis. The effector caspase zymogens are unique among the caspase subfamilies in that the proteins are stable, yet inactive, dimers (13). In addition to differences in oligomeric states, the caspase-hemoglobinase fold is also the basis for the evolution of separate enzyme specificity and allosteric regulation (31, 32). However, the evolutionary processes that resulted in enzymatic, allosteric, and oligomeric diversity of the caspases are unknown. Effector caspases-3, −6, and −7 have a significant role in apoptosis and serve overlapping but nonredundant functions (33), and multiple studies have examined enzyme specificity and regulation of extant caspases (20, 31, 34, 35). In addition, we have previously examined evolutionary changes resulting in amino acid substitutions that affect enzyme specificity (15, 36) and allosteric regulation (25, 37, 38), but there is a dearth of information regarding changes in the caspase folding landscape. To date, only human caspase-3 has been examined in detail (6, 26, 27, 39), so it was not clear whether all effector caspases utilized the same folding landscape.

Here, we examined the conservation of the folding landscape by determining the equilibrium folding/unfolding process of the common ancestor of effector caspases as well as human procaspases-6 and −7, and we compared the results to our previous data for human procaspase-3. Collectively, the data provide a baseline for understanding the folding landscape of effector caspases and serve as a platform for examining the monomeric caspase subfamilies as well as evolutionary changes that resulted in the stable dimer. The PCP3 dimer folds and assembles through two partially folded intermediate conformations, a monomer (I) and a dimer (I_2_), with an overall conformational free energy, ΔG°_conf_, of ~22 kcal/mol at pH >6 (26, 27). Our data for the folding and assembly of the common ancestor of effector caspases, PCP-CA, show that the monomeric intermediate (I) is present in the folding pathway, but we do not observe the dimeric intermediate (I_2_). In contrast, our data for the folding and assembly of the PCP6 dimer show that the dimeric intermediate (I_2_) is present in the folding pathway, but we do not observe the monomeric intermediate (I). Finally, PCP7 is more similar to the common ancestor, with the monomeric intermediate (I) present in the folding pathway. We suggest that a parsimonious explanation for the data is that the effector caspase folding landscape consists of both intermediates, I and I_2_, and that the population distribution of I relative to I_2_ varies among the three extant caspases. In this case, the more prevalent intermediate, I or I_2_, is observed in our spectroscopic assays. For PCP3, both intermediates are observed due to their similar stability, resulting in a substantial population of both species. We note that our data do not rule out the possibility that the dimeric intermediate arose separately in PCP3 and in PCP6 after the three lineages split ~450 million years ago. Nevertheless, such a folding landscape, where both intermediates are accessible for evolutionary changes, would provide flexibility in a protein fold that is utilized by multiple subfamilies. In this case, amino acid changes that occur through evolution could stabilize one intermediate relative to the other in a species-dependent manner to provide for differences in cellular and environmental conditions of the organism. Our equilibrium data presented here do not provide information on the rate of folding, so further kinetic studies would determine whether changes in the relative distribution of I and I_2_ affect folding efficiency. We note, however, that mutations in PCP3 that affect the rate of I-to-I_2_ dimerization did result in formation of a dimerization-incompetent monomer, leading to hysteresis in the equilibrium folding data (39).

Although the folding landscape of caspases has been conserved for >650 million years, amino acid substitutions through evolution have resulted in differences in stability among the extant caspases. When comparing the overall conformational stability of the extant human effector caspases, one observes that the dimer generally falls in the range of ~17-30 kcal/mol (Supplemental Table S2). PCP6 is the most stable, with ΔG°_conf_, of ~30 kcal/mol at pH >6, and PCP7 is the least stable, with ΔG°_conf_, of ~17 kcal/mol at pH >6. PCP3 and PCP-CA show similar stabilities, with ΔG°_conf_, of ~22 kcal/mol at pH >6. The unfolding data also show that the dimer of PCP6 is more stable than that of PCP7 or of PCP-CA, where the mid-point of dimer dissociation is approximately 5 M urea for PCP6 *versus* 2 M urea for PCP7 and PCP-CA, likely due to the higher population of the I_2_ intermediate in PCP6 compared to PCP7 and PCP-CA. Notably, Matthews and others (40, 41) have reported that amino acid substitutions that change in the number of salt bridges in the native structure of a protein are a key parameter in modulating the free energy landscape of a protein fold. We compared the number of salt bridges among extant caspases using a protein tool developed by Ferruz et al (42). The analysis showed that PCP3 (PDB:1CP3), PCP6 (PDB:3S70), and PCP7 (PDB:1F1J) consist of 21, 22, and 15 salt bridges, respectively. Although the data are consistent with PCP7 exhibiting the lowest conformational free energy, further examination of the intersubunit contacts would provide a quantitative assessment of the contributions to overall stability.

Our data show that the conformational stability is similar for the different species in the folding landscape, so based on the data for the four effector caspases, we compared the ΔG°_conf_ for each transition to determine a range of conformational free energy for the native dimer (N_2_), the dimeric intermediate (I_2_), and the monomeric intermediate (I) (Figure 7). We used ΔG°_conf_ values at the higher pHs since three of the caspases (PCP3, PCP7, and PCP-CA) were less stable at low pH. Essentially, the values of 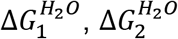, and 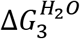 from Supplemental Table S2 were utilized in the analysis. The comparison shows that the native dimer of effector caspases has a ΔG°_conf_, of ~6.7 kcal/mol with a range of 4.9-8.7 kcal/mol. The dimeric intermediate is more stable, with ΔG°_conf_, of ~11.3 kcal/mol, within a range of 9.5-15 kcal/mol, and the monomeric intermediate is similar to that of the dimer, with ΔG°_conf_, of ~6.4 kcal/mol, within a range of 5.2-8.5 kcal/mol. Combining the three species results in the overall conformational free energy of 25.7 kcal/mol, with a broad range of 15-34 kcal/mol. We suggest that the folding landscape of effector caspases was established in the common ancestor and provides two partially folded conformations, one monomer and one dimer, from which evolutionary changes can establish the relative distribution of the intermediates as well as the overall conformational stability of the dimer, within the ranges shown in Figure 7.

**Figure 7:**
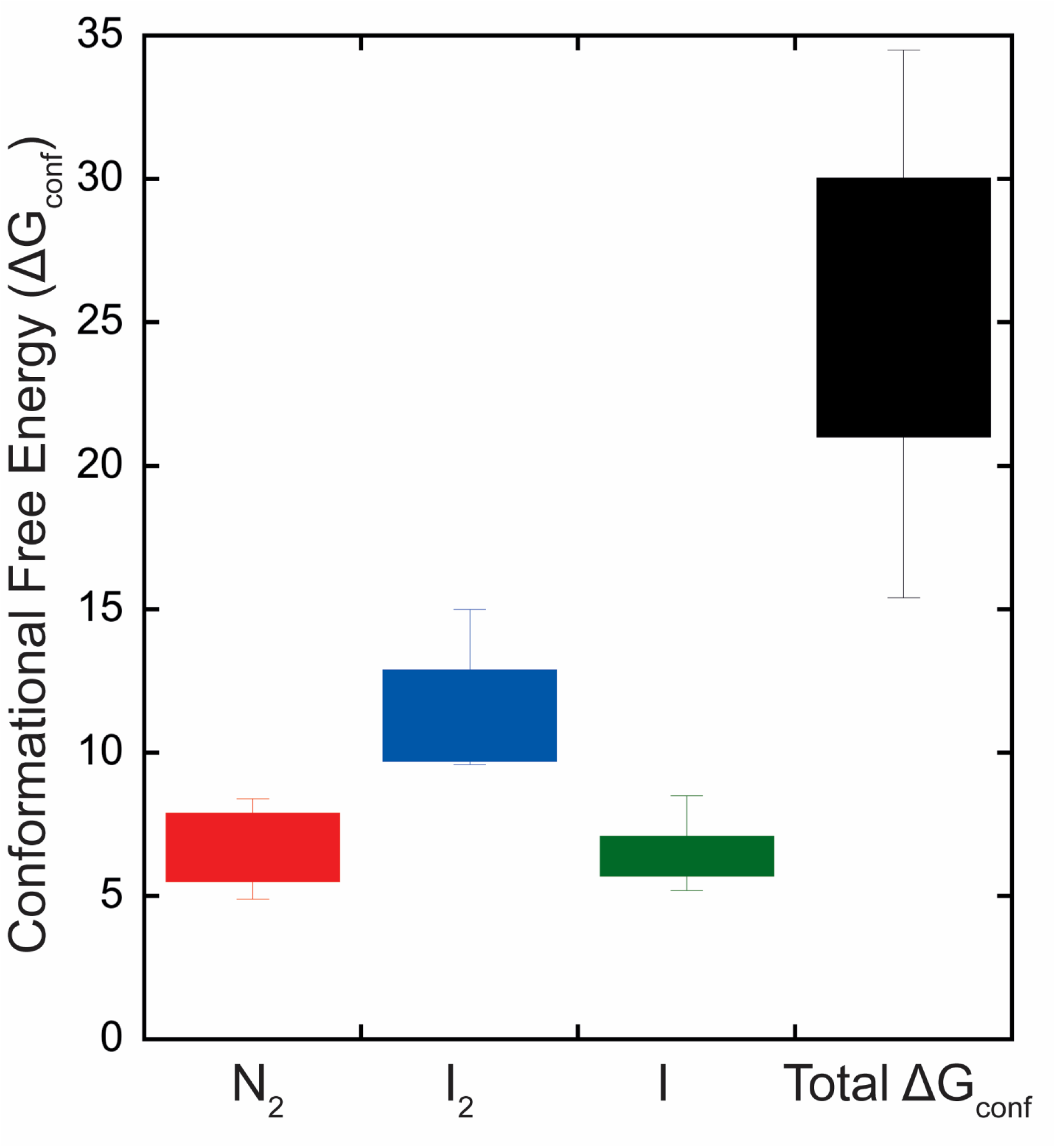
Conformational free energies of three conformations (N_2_, I_2_, and I) in the caspase folding landscape as well as the total conformational free energy for effector caspases. Values were calculated as described the text using data in Supplemental Table S2.

We showed previously that changes in pH are an excellent perturbation of the caspase folding landscape because the PCP3 dimer dissociates below pH 5 such that the protein is a monomer at pH 4 (27). Furthermore, we suggested that two histidine residues in PCP3 stabilize the dimer at higher pH through charge-charge interactions across the dimer interface. In contrast to PCP3, PCP6 remains dimeric over the entire pH range examined here (pH 4-8.5), and we note that CP-H186 is unique in PCP3 (see Figure 2). At the CP-186 position, one observes glutamate in PCP6 and in PCP-CA, and glutamine in PCP7. In addition, the PCP7 dimer is less stable at lower pH, but the protein remains dimeric. At present, it is not clear why the PCP-CA is less stable at lower pH, but the data suggest that evolutionary changes stabilized the effector caspase dimer against pH changes, except for PCP3. We noted previously (27) that pH changes in the cell during apoptosis (43, 44) may affect the monomer-dimer equilibrium of caspase-3, resulting in lower activity overall for the pool of caspase-3 in the apoptotic cell, since the monomer is enzymatically inactive. In this case, the pool of caspase-6 and/or caspase-7 activity would be affected only by the protonation/deprotonation of the catalytic cysteine-histidine residues rather than the additional regulatory feature of monomer-dimer equilibrium. Since PCP7 and PCP-CA were unfolded irreversibly around pH 5 and 6, it is not clear at what pH the protein dissociates to monomers or why the dimer is less stable at lower pH. Overall, however, the differences in pH effects suggest that caspase-6 could be the main executioner caspase in a low pH environment rather than caspase-3. Caspase-6 has a more specialized role in specific physiological contexts compared to caspases-3 and −7 (45–47). Further analysis of the differences in the dimeric interface among effector caspases would provide clarity regarding the evolutionary changes that resulted in the higher conformational stability of the PCP6 dimer at low pH.

In summary, by comparing the equilibrium folding/unfolding of the human effector caspases and their common ancestor, we show that the folding landscape was established in the common ancestor (>650 Mya) and that evolution can utilize intermediates in the landscape to effect changes in the conformational stability of extant caspases, within the constraints of the caspase-hemoglobinase fold. The caspase family highlights how sequence changes through evolutionary processes can provide species-dependent flexibility in the caspase dimer. Hence, the stability of the native enzyme and the response to changes in the environment can be fine-tuned in a species-specific manner while retaining the overall caspase-hemoglobinase fold.

## Experimental procedures

### Cloning, protein expression, and protein purification

For all procaspase proteins, the catalytic cysteine (CP-117, Figure 2) was mutated to serine using site-directed mutagenesis, as described previously (48). The inactive procaspases were cloned into the pET11a expression vector with a C-terminal hexahistidine tag, and all proteins were expressed in *E. coli* BL21(DE3) pLysS cells and purified as previously described (30, 49, 50).

### Sample preparation for equilibrium unfolding

Denaturation and renaturation experiments were carried out as described previously (51). Briefly, urea stock solutions (10 M) were prepared in citrate buffer (20 mM sodium citrate/citric acid, pH 4 to pH 5.5, 1 mM DTT), phosphate buffer (20 mM potassium phosphate monobasic and dibasic, pH 6 to 8, 1 mM DTT), or Tris buffer (20 mM Tris-HCl, pH 8.5, 1 mM DTT). For unfolding experiments, samples were prepared in the corresponding buffer with final urea concentrations between 0 and 8 M. Stock protein in buffer was added such that the final concentrations are as shown in the figures. For renaturation experiments, the protein was first incubated in an 8 M urea-containing buffer for 3 hours at 25 °C. The unfolded protein was then diluted with the corresponding buffer and urea such that the final urea concentrations were between 0.5 and 8 M. For all equilibrium unfolding experiments, protein concentrations from 0.5 to 4 μM were used. In both denaturation and renaturation experiments, the samples were incubated at 25 °C for a minimum of 16 h to allow for equilibration. This incubation time was found to be optimal to allow the protein to reach equilibrium at all urea concentrations.

### Fluorescence emission and circular dichroism (CD) measurements

Fluorescence emission was acquired using a PTI C-61 spectrofluorometer (Photon Technology International, Birmingham, NJ) from 300 nm to 400 nm following excitation at 280 or 295 nm. Excitation at 280 nm follows tyrosinyl and tryptophanyl fluorescence emission, whereas excitation at 295 nm follows the tryptophanyl fluorescence emission. CD measurements were recorded using a J-1500 CD spectropolarimeter (Jasco) between 220 and 240 nm. Fluorescence and CD spectra were measured using a 1 cm path length cuvette and constant temperature (25 °C). All data were corrected for buffer background.

### Data analysis and global fits of the equilibrium unfolding data

The data were fit globally and interpreted as described previously (26, 27, 51). Briefly, fluorescence emission and CD data were collected between pH 8.5 and 4 for all three proteins and at three to four protein concentrations, which resulted in 9 to 12 data sets at each pH. The data were fit to a 2-state or 3-state equilibrium folding model, as described below. At pH 7.5, the data for PCP6 were best fit to a 3-state equilibrium folding model described with a dimeric intermediate in equilibrium with the native and unfolded protein, as shown in eq 1. In contrast, the data for PCP7 and PCP-CA were best fit to a 3-state equilibrium folding model described with a monomeric intermediate in equilibrium with the native and unfolded protein, as shown in eq 2.

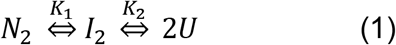

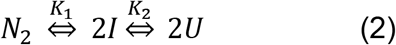

In both equations 1 and 2, K_1_ and K_2_ refer to equilibrium constants for the two steps, respectively. At pH 4, the data for PCP6 was best fit to a two-state equilibrium folding model, where the native dimer is in equilibrium with the unfolded protein, as shown in equation 3.

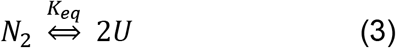

In contrast to PCP6, at pH 4 the data for PCP7 was best fit to a 3-state equilibrium folding model for a monomer, as shown in equation 4.

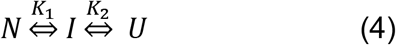

For all proteins, the equilibrium folding/unfolding data at each pH were fit globally using the appropriate folding model from equations 1-5 and the program Igor Pro (WaveMetrics, Inc.), as described previously (26, 27, 51). Results from the fits are shown in Table 1 and as the solid lines in Figure 4 and Supplemental Figures S2-S6. The change in solvent accessible surface area was calculated as describe by Scholtz and co-workers (29), as shown in equation 5.

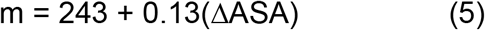

## Data Availability

All data are contained in the manuscript and in Supporting Information.

## Supporting Information

This article contains supporting information.

## Funding

This work was supported by a grant from the National Institutes of Health [grant number GM127654 (to A.C.C)]. The content is solely the responsibility of the authors and does not necessarily represent the official views of the National Institutes of Health.

## Conflict of interest

The authors declare that they have no conflicts of interest with the contents of this article.

## Notes

### Competing Interest Statement

The authors have declared no competing interest.

