## Supplemental Figures Tables for "Evolution of the folding landscape of effector caspases"

Running Title: Common intermediates in the caspase folding landscape

Keywords: caspase, protein folding, folding landscape, protein evolution, oligomerization, evolutionary biology, evolution, apoptosis, protease,

### Supporting Information

**Supplemental Table S1**: Amino acids composition and theoretical parameters of procaspase-3, -6, -7, and their common ancestor (PCP-CA).

| **Composition** [(1)] | **Proteins** | | | |
|---|---|---|---|---|
| | PCP3 | PCP6 | PCP7 | PCP-CA |
| **Total Number Amino Acids** | 277 | 293 | 303 | 275 |
| **Ala (A)** | 12 (4.3%) | 19 (6.5%) | 18 (5.9%) | 9 (3.3%) |
| **Arg (R)** | 14 (5.1%) | 17 (5.8%) | 15 (5.0%) | 7 (2.5%) |
| **Asn (N)** | 15 (5.4%) | 11 (3.8%) | 14 (4.6%) | 13 (4.7%) |
| **Asp (D)** | 20 (7.2%) | 20 (6.8%) | 27 (8.9%) | 20 (7.3%) |
| **Cys (C)** | 8 (2.9%) | 10 (3.4%) | 11 (3.6%) | 7 (2.5%) |
| **Gln (Q)** | 4 (1.4%) | 7 (2.4%) | 11 (3.6%) | 10 (3.6%) |
| **Glu (E)** | 20 (7.2%) | 20 (6.8%) | 19 (6.3%) | 25 (9.1%) |
| **Gly (G)** | 16 (5.8%) | 19 (6.5%) | 18 (5.9%) | 18 (6.5%) |
| **His (H)** | 8 (2.9%) | 12 (4.1%) | 7 (2.3%) | 6 (2.2%) |
| **Ile (I)** | 19 (6.9%) | 13 (4.4%) | 17 (5.6%) | 14 (5.1%) |
| **Leu (L)** | 20 (7.2%) | 26 (8.9%) | 20 (6.6%) | 25 (9.1%) |
| **Lys (K)** | 22 (7.9%) | 20 (6.8%) | 25 (8.3%) | 28 (10.2%) |
| **Met (M)** | 10 (3.6%) | 7 (2.4%) | 7 (2.3%) | 6 (2.2%) |
| **Phe (F)** | 15 (5.4%) | 18 (6.1%) | 17 (5.6%) | 12 (4.4%) |
| **Pro (P)** | 7 (2.5%) | 10 (3.4%) | 12 (4.0%) | 9 (3.3%) |
| **Ser (S)** | 26 (9.4%) | 18 (6.1%) | 21 (6.9%) | 26 (9.5%) |
| **Thr (T)** | 16 (5.8%) | 16 (5.5%) | 15 (5.0%) | 13 (4.7%) |
| **Trp (W)** | 2 (0.7%) | 2 (0.7%) | 2 (0.7%) | 1 (0.4%) |
| **Tyr (Y)** | 10 (3.6%) | 10 (3.4%) | 9 (3.0%) | 13 (4.7%) |
| **Val (V)** | 13 (4.7%) | 18 (6.1%) | 18 (5.9%) | 13 (4.7%) |
| **Molecular Weight (Da)** | 31,608 | 33,310 | 34,277 | 31221 |
| **pI** | 6.09 | 6.46 | 5.72 | 5.2 |
| **Extinction Coefficient (280 nm, M-1cm-1)** [(2)] | 26,500 | 25,900 | 24,410 | 24870 |

[1] Parameters exclude the LEHHHHHH C-terminal tag.

[2] Assuming all cysteine residues are reduced.

**Supplemental Table S1**: Free energy changes of each step of folding/unfolding of extant and ancestral effector caspases.

| pH | PCP6 | | | PCP7 | | | PCP-CA | | | PCP3* | | | |
|---|---|---|---|---|---|---|---|---|---|---|---|---|---|
| | $\Delta G_1^{H_2O}$ | $\Delta G_2^{H_2O}$ | $\Delta G_{total}^{H_2O}$ | $\Delta G_1^{H_2O}$ | $\Delta G_2^{H_2O}$ | $\Delta G_{total}^{H_2O}$ | $\Delta G_1^{H_2O}$ | $\Delta G_2^{H_2O}$ | $\Delta G_{total}^{H_2O}$ | $\Delta G_1^{H_2O}$ | $\Delta G_2^{H_2O}$ | $\Delta G_3^{H_2O}$ | $\Delta G_{total}^{H_2O}$ |
| 4 | - | 27.6±0.8 | 27.6 | 6.9±0.5 | 5.2±0.3 | 12.1 | 0.4±0.2 | 5.1±0.3 | 5.5 | - | - | 3.8±0.2 | 3.8±0.2 |
| 4.2 | | | | | | | | | | - | 1.3±0.3 | 3.7±0.3 | 5±0.6 |
| 4.5 | 0.9±0.1 | 30.4±1.4 | 31.3 | - | - | - | - | - | - | - | 1.7±0.6 | 5.3±0.5 | 7±1.1 |
| 4.75 | | | | | | | | | | 0.8±0.2 | 9.2±0.1 | 5±0.2 | 15±0.5 |
| 5 | 0.4±0.5 | 29.5±1 | 29.9 | - | - | - | - | - | - | 3.1±0.6 | 9.6±0.2 | 5.3±0.3 | 18.1±1 |
| 5.5 | 0.3±0.3 | 28.5±1.4 | 28.8 | - | - | - | - | - | - | 3.3±0.1 | 11±1.9 | 6.8±1.8 | 21.1±3.8 |
| 6 | 5.3±0.9 | 26.1±0.9 | 31.4 | - | - | - | - | - | - | 6.1±1.0 | 10.3±0.3 | 5.6±0.2 | 22±1.5 |
| 6.5 | 6±0.9 | 27.6±1 | 33.6 | - | - | - | 12.8±0.4 | 8.5±0.5 | 21.3 | 5.9±0.8 | 10.7±0.6 | 5.9±0.6 | 22.6±2 |
| 7 | 5.5±0.7 | 23.8±0.2 | 29.3 | - | - | - | - | - | - | 8.3±1.3 | 10.5±1 | 7±0.5 | 25.8±2.8 |
| 7.5 | 8.4±0.8 | 24.4±0.9 | 32.8 | 10.2±0.2 | 5.2±0.1 | 15.4 | 14.9±0.1 | 6.9±0.3 | 21.8 | 7.9±0.1 | 9.7±0.3 | 7.2±0.5 | 24.8±0.9 |
| 8 | 6.8±0.6 | 27.7±1.6 | 34.5 | 12.9±0.1 | 5.5±1.3 | 18.4 | - | - | - | 5.8±0.8 | 9.6±0.2 | 7.8±0.4 | 23.2±1.4 |
| 8.5 | 5.4±0.5 | 25.3±1.3 | 30.8 | - | - | - | 13.8±0.8 | 5.8±0.3 | 19.6 | 4.9±0.6 | 9.6±0.3 | 6.2±0.6 | 20.7±1.5 |

* Data from Bose and Clark (1, 2).

**Supplemental Table S3**: Cooperative index (m-values) of each step of folding/unfolding of extant and ancestral effector caspases.

| pH | PCP6 | | | PCP7 | | | PCP-CA | | | PCP3* | | | |
|---|---|---|---|---|---|---|---|---|---|---|---|---|---|
| | $m_1$ | $m_2$ | $m_{total}$ | $m_1$ | $m_2$ | $m_{total}$ | $m_1$ | $m_2$ | $m_{total}$ | $m_1$ | $m_2$ | $m_2$ | $m_{total}$ |
| 4 | - | 3.9±0.2 | 3.9 | 1.0±0.1 | 1.3±0.2 | 2.3 | 0.7±0.1 | 1.1±0.1 | 1.8 | - | - | 1.1±0.1 | 1.1±0.1 |
| 4.2 | | | | | | | | | | - | 1.5±0.2 | 0.9±0.1 | 2.4±0.3 |
| 4.5 | 0.9±0.1 | 3.5±0.2 | 4.4 | - | - | - | - | - | - | - | 2.2±0.4 | 1.2±0.1 | 3.4±0.5 |
| 4.75 | | | | | | | | | | 2.8±0.1 | 0.9±0.2 | 1.2±0.1 | 4.9±0.4 |
| 5 | 0.8±0.1 | 3.2±0.2 | 4 | - | - | - | - | - | - | 2.3±0.3 | 0.2±0.1 | 1.2±0.1 | 3.7±0.5 |
| 5.5 | 0.7±0.1 | 3.2±0.2 | 3.9 | - | - | - | - | - | - | 3±0.1 | 0.4±0.2 | 1.5±0.2 | 4.9±0.5 |
| 6 | 2.7±0.4 | 3.1±0.2 | 5.8 | - | - | - | - | - | - | 3.4±0.5 | 0.2±0.1 | 1.3±0.1 | 4.9±0.7 |
| 6.5 | 2.4±0.3 | 3.4±0.2 | 5.8 | - | - | - | 1.5±0.1 | 1.5±0.1 | 3 | 2.6±0.4 | 0.4±0.1 | 1.3±0.1 | 4.3±0.6 |
| 7 | 2±0.2 | 2.8±0.2 | 4.8 | - | - | - | - | - | - | 2.8±0.5 | 0.5±0.1 | 1.2±0.1 | 4.5±0.7 |
| 7.5 | 2.6±0.3 | 2.9±0.2 | 5.5 | 1.3±0.1 | 1.2±0.1 | 2.5 | 2±0.1 | 1.2±0.1 | 3.2 | 2.8±0.1 | 0.4±0.1 | 1.2±0.1 | 4.4±0.3 |
| 8 | 2.2±0.2 | 3.5±0.3 | 5.7 | 1.6±0.1 | 1.3±0.1 | 2.9 | - | - | - | 2.8±0.4 | 0.4±0.1 | 1.5±0.1 | 4.7±0.6 |
| 8.5 | 1.9±0.2 | 3.4±0.3 | 5.3 | - | - | - | 2±0.2 | 1±0.1 | 3 | 1.9±0.2 | 0.4±0.1 | 1±0.1 | 3.3±0.4 |

* Data from Bose and Clark (1, 2).

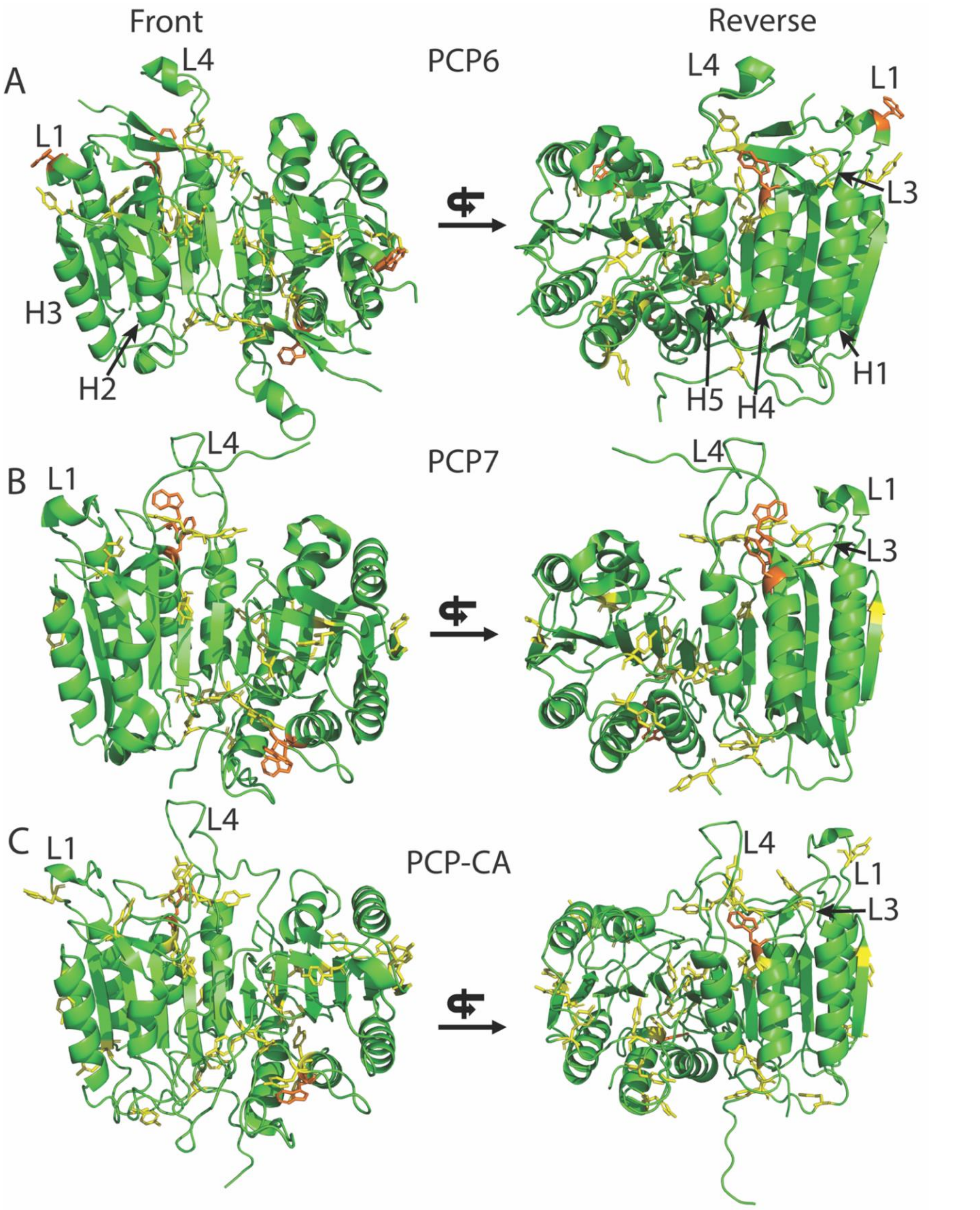


**Supplemental Figure S1**. Tyrosine residues (yellow sticks) and tryptophan residues (orange sticks) are shown in PCP6 (PDB:4nbk) (**A**), PCP7 (PDB:1K86) (**B**) and PCP-CA (model structure) (**C**). Helices H1-H5 are labeled. L1, L3, and L4 refer to active site loops 1, 3, and 4, respectively.

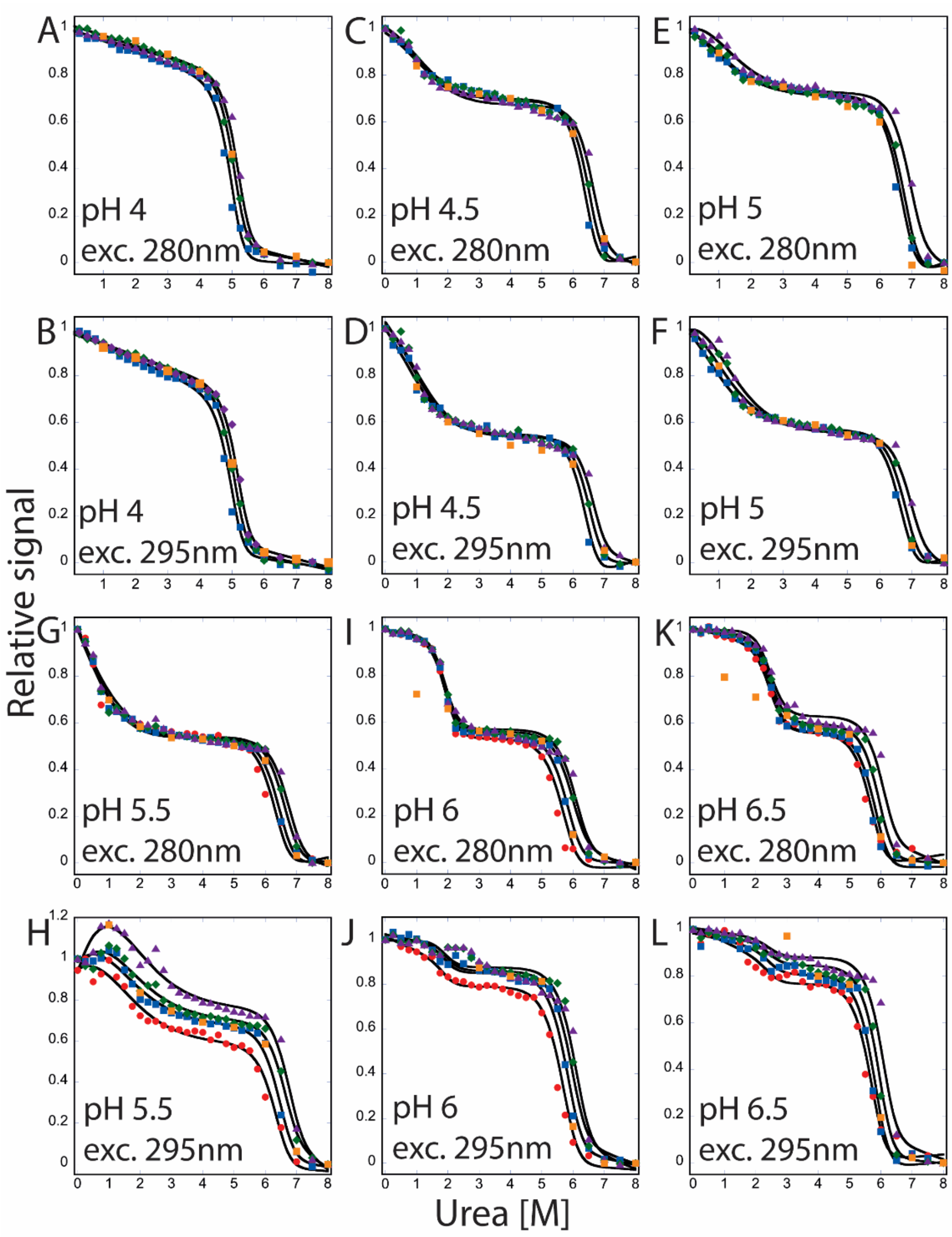


**Supplemental Figure S2**: Equilibrium unfolding monitored by fluorescence emission for PCP6 from pH 4 to 6.5. Relative signal of fluorescence emission with excitation at 280 nm and 295 nm at pH 4 (**A, B**), pH 4.5 (**C, D**), pH 5 (**E, F**), pH 5.5 (**G, H**), pH 6 (**I, J**), and pH 6.5 (**K, L**). Three different protein concentrations were used from pH 4, 4.5 and 5, and four different protein concentrations were used from pH 5.5, 6 and 6.5. Colored solid symbols represent raw data and corresponding solid lines represent the global fits of the data in an appropriate model described in text. The following protein concentrations were used: 0.5 μM (●), 1 μM (■), 2 μM (◆), and 4 μM (▲). Orange squares (■) represent refolding data at 2 μM protein to show reversibility.

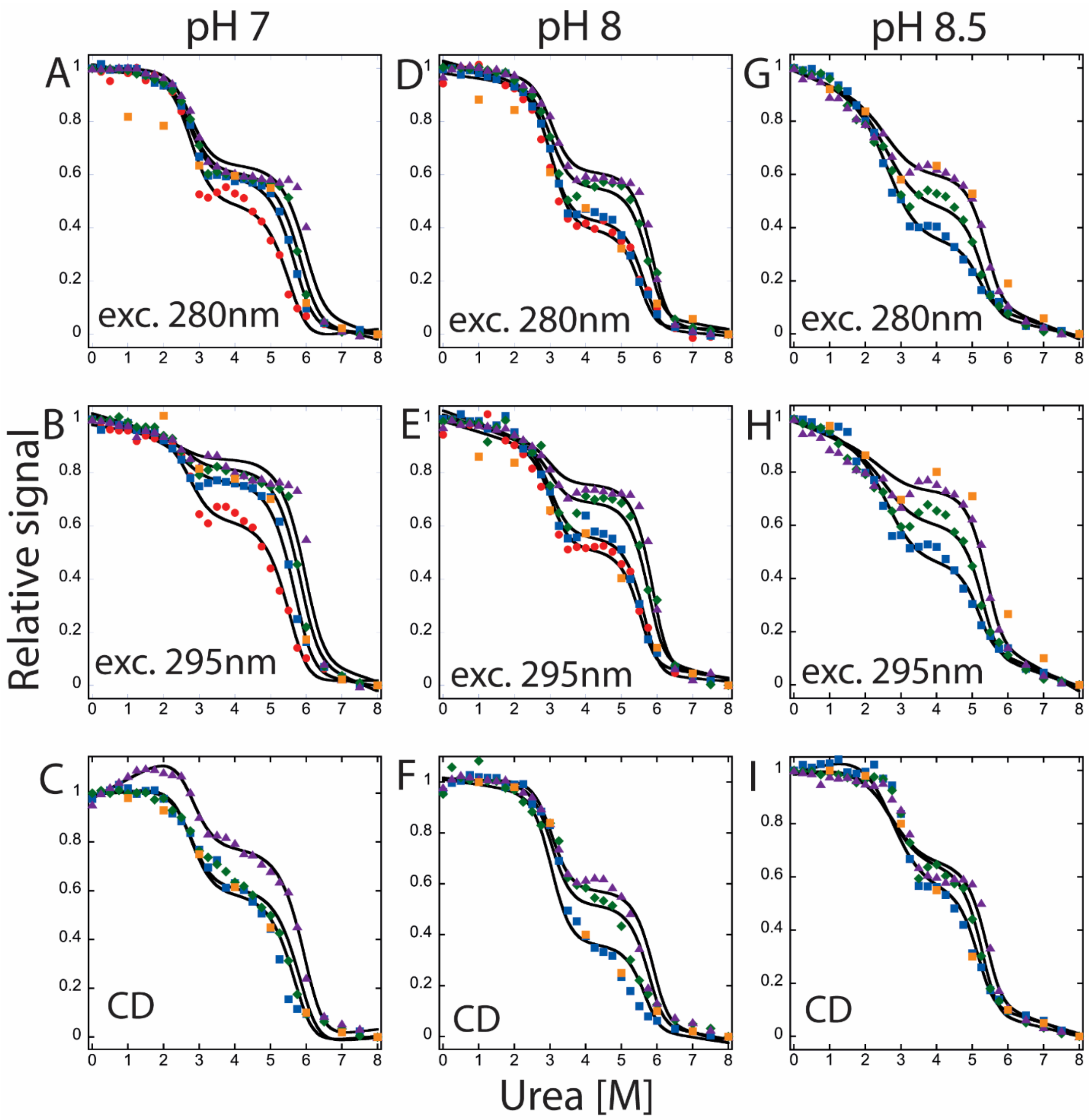


**Supplemental Figure S3**: Equilibrium unfolding of PCP6 from pH 7 to 8.5. Relative signal of fluorescence emission with excitation at 280 nm, 295 nm, and CD at pH 7 (**A, B, C**), pH 8 (**D, E, F**), and pH 8.5 (**G, H, I**). Colored solid symbols represent raw data and corresponding solid lines represent the global fits of the data in an appropriate model described in text. Symbols are represented as follows. 0.5 µM (●), 1 µM (■), 2 µM (◆), and 4 µM (▲). Orange squares (■) represent refolding data at 2 µM protein to show reversibility.

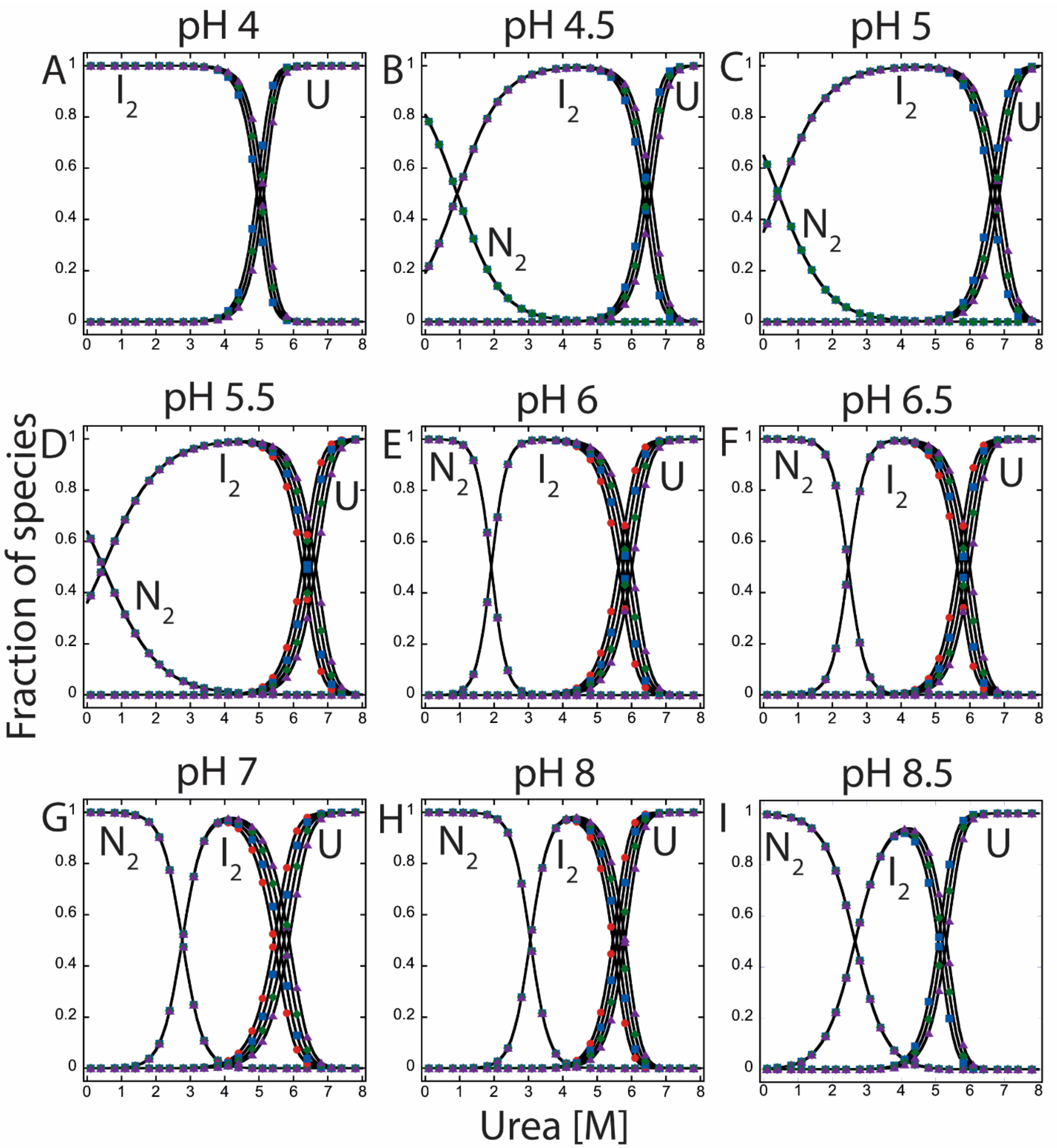


**Supplemental Figure S4**: Fraction of species of PCP6 folding/unfolding as a function of urea concentration from pH 4 to 8.5. The fractions of native, intermediate, and unfolded protein were calculated as a function of urea concentration from fits of the data at the respective pH. $N_2$ refers to dimeric native protein, $I_2$ is the dimeric intermediate, and U refers to the unfolded species. The following protein concentrations were used in the calculations: 0.5 µM (●), 1 µM (■), 2 µM (◆), and 4 µM (▲).

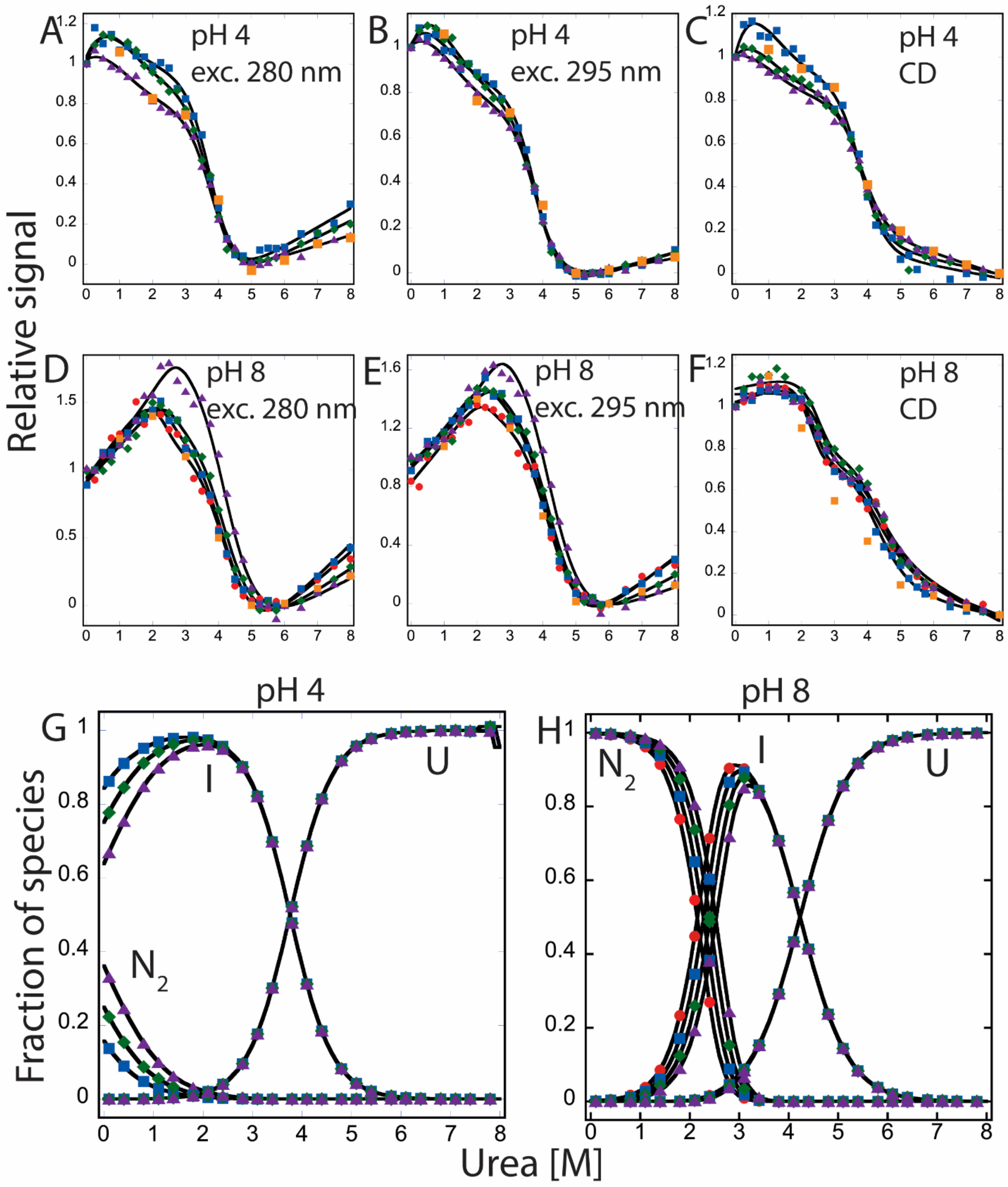


**Supplemental Figure S5**: Fits of equilibrium unfolding data and fraction of species of PCP7. Relative signal of fluorescence emission with excitation at 280 nm and 295 nm, and CD at pH 4 (**A, B, C**), and pH 8 (**D, E, F**), and calculated fraction of species of pH 4 (**G**), and pH 8 (**H**). $N_2$ refers to dimeric native protein, I is monomeric intermediate, and U refers to unfolded species. The following protein concentrations were used: 0.5 µM (●), 1 µM (■), 2 µM (◆), and 4 µM (▲). Orange squares (■) represent refolding data of 2 µM protein to show reversibility.

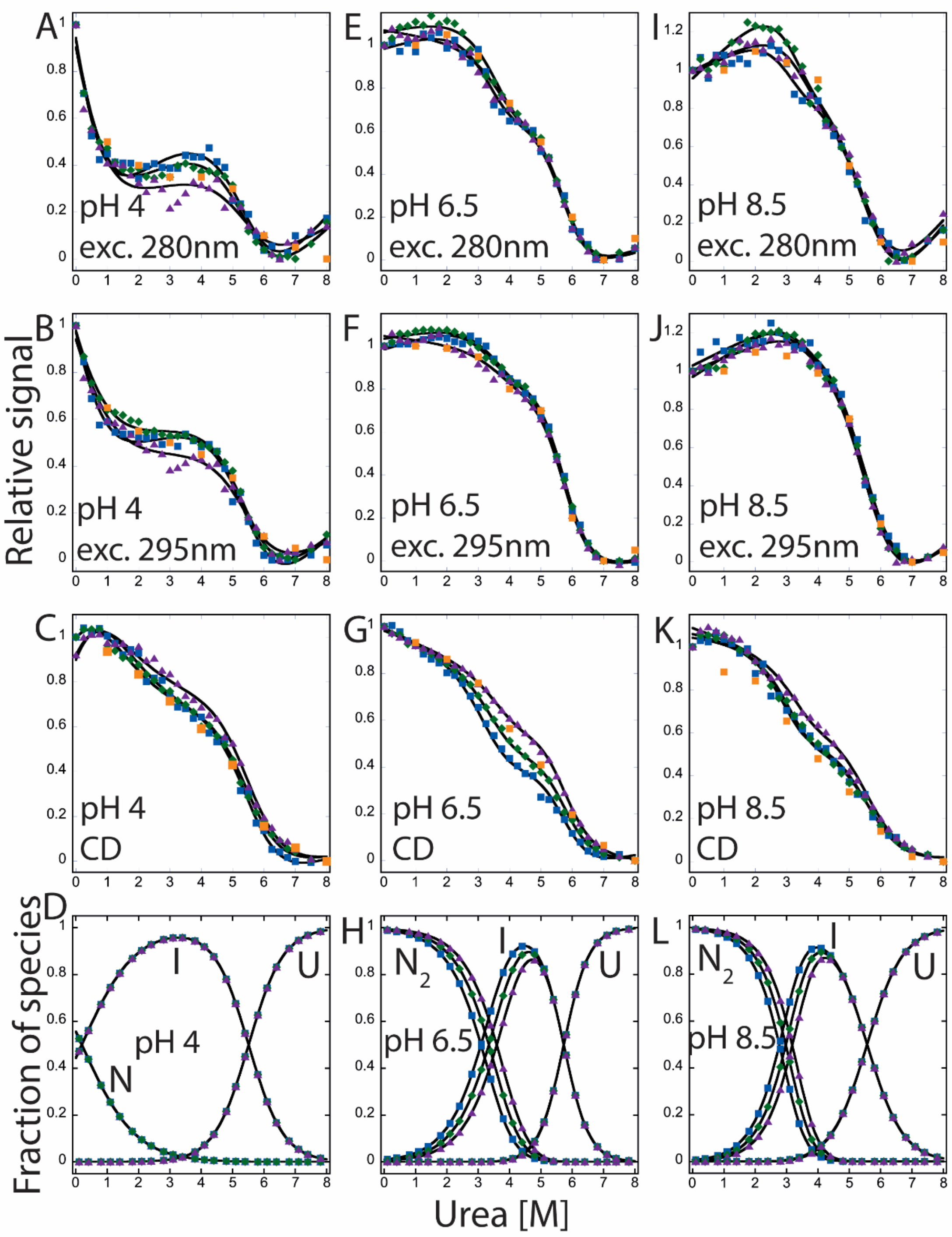


**Supplemental Figure S6**: Fits of equilibrium unfolding data and fraction of species of PCP-CA. Relative signal of fluorescence emission with excitation at 280 nm and 295 nm, and CD at pH 4 (**A, B, C**), pH 6.5 (**E, F, G**), and pH 8.5 (**I, J, K**) and calculated fraction of species at pH 4 (**D**), pH 6.5 (**H**), and pH 8.5 (**L**). $N_2$ refers to dimeric native protein, N is monomeric "native" protein, I is monomeric intermediate, and U refers to unfolded species. The following protein concentrations were used: 1 µM (■), 2 µM (◆), and 4 µM (▲). Orange squares (■) represent refolding data of 2 µM protein to show reversibility.

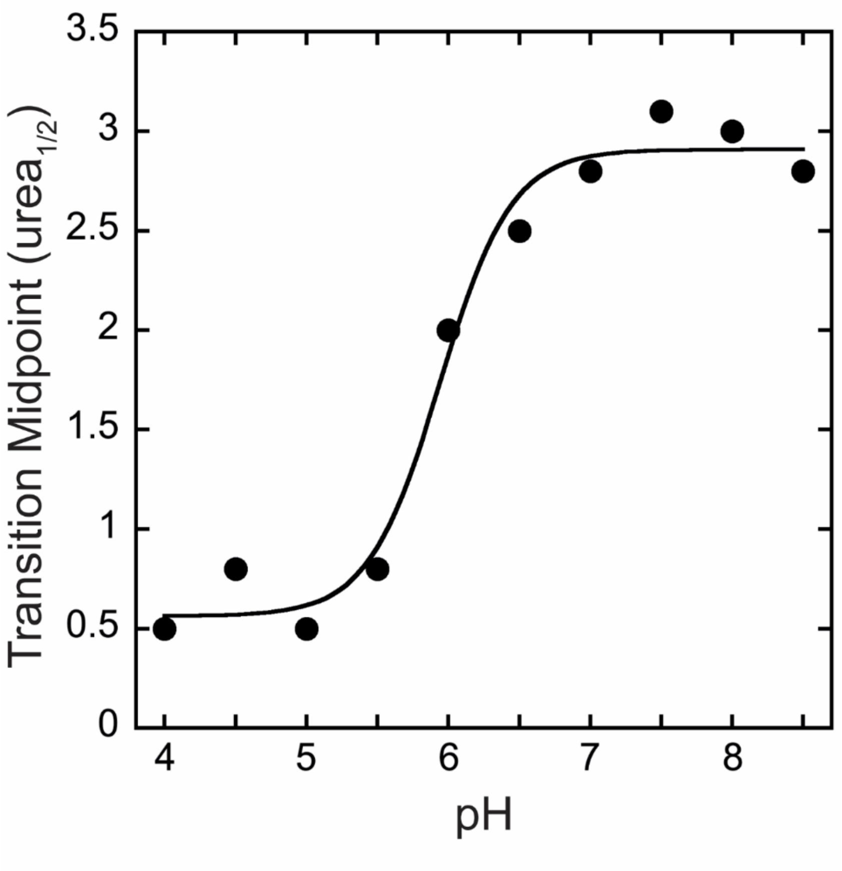


**Supplemental Figure S7**: Changes in the mid-point of the first transition ($N_2 \rightleftarrows I_2$) in unfolding of PCP6 over the pH range of 4 to 8.5. The solid line represents fits of the data as described previously (3). The pKa of the transition was determined to be 5.9.
